# *Toxoplasma gondii* harbors a hypoxia-responsive coproporphyrinogen dehydrogenase-like protein

**DOI:** 10.1101/2023.11.16.567449

**Authors:** Melanie Key, Carlos Gustavo Baptista, Amy Bergmann, Katherine Floyd, Ira J. Blader, Zhicheng Dou

## Abstract

*Toxoplasma gondii* is an apicomplexan parasite that is the cause of toxoplasmosis, a potentially lethal disease for immunocompromised individuals. During *in vivo* infection, the parasites encounter various growth environments, such as hypoxia. Therefore, the metabolic enzymes in the parasites must adapt to such changes to fulfill their nutritional requirements. *Toxoplasma* can *de novo* biosynthesize some nutrients, such as heme. The parasites heavily rely on their own heme production for intracellular survival. Notably, the antepenultimate step within this pathway is facilitated by coproporphyrinogen III oxidase (CPOX), which employs oxygen to convert coproporphyrinogen III to protoporphyrinogen IX through oxidative decarboxylation. Conversely, some bacteria can accomplish this conversion independently of oxygen through coproporphyrinogen dehydrogenase (CPDH). Genome analysis found a CPDH ortholog in *Toxoplasma*. The mutant *Toxoplasma* lacking CPOX displays significantly reduced growth, implying that TgCPDH potentially functions as an alternative enzyme to perform the same reaction as CPOX under low oxygen conditions. In this study, we demonstrated that TgCPDH exhibits coproporphyrinogen dehydrogenase activity by complementing it in a heme synthesis-deficient *Salmonella* mutant. Additionally, we observed an increase in TgCPDH expression in *Toxoplasma* when it grew under hypoxic conditions. However, deleting *TgCPDH* in both wildtype and heme-deficient parasites did not alter their intracellular growth under both ambient and low oxygen conditions. This research marks the first report of a coproporphyrinogen dehydrogenase-like protein in eukaryotic cells. Although TgCPDH responds to hypoxic conditions and possesses enzymatic activity, our findings suggest that it does not directly affect intracellular infection or the pathogenesis of *Toxoplasma* parasites.

**IMPORTANCE:** *Toxoplasma gondii* is a ubiquitous parasite capable of infecting a wide range of warm-blooded hosts, including humans. During its lifecycle, these parasites must adapt to varying environmental conditions, including situations with low oxygen levels. Our research, in conjunction with studies conducted by other laboratories, has revealed that *Toxoplasma* primarily relies on its own heme production during acute infections. Intriguingly, in addition to this classical heme biosynthetic pathway, the parasites encode a putative oxygen-independent coproporphyrinogen dehydrogenase, suggesting its potential contribution to heme production under varying oxygen conditions, a feature typically observed in simpler organisms like bacteria. Notably, so far, coproporphyrinogen dehydrogenase has only been identified in some bacteria for heme biosynthesis. Our study discovered that *Toxoplasma* harbors a functional enzyme displaying coproporphyrinogen dehydrogenase activity, which alters its expression in the parasites when they face fluctuating oxygen levels in their surroundings.

## INTRODUCTION

Infectious pathogens encounter diverse oxygen conditions throughout their life cycles. For example, pathogenic bacteria inhabiting the host’s intestine are exposed to a notably reduced oxygen environment, ranging from 5% -0.5% (1) compared to 21% ambient conditions. This substantial fluctuation in oxygen levels demands an adaptive response from these pathogens to effectively cope with hypoxic conditions. Similarly, *Toxoplasma gondii* needs to deal with a substantial reduction in oxygen while inhabiting the host’s organs during *in vivo* infection. To adapt to hypoxic conditions, the parasites activate the host’s hypoxia-inducible factor (HIF) to stabilize the HIF1-α subunit to upregulate host hexokinase 2 expression and activity (2). This is achieved by inhibiting the activity of host prolyl hydroxylase 2, an α-ketoglutarate-dependent dioxygenase whose high K_m_ towards oxygen allows it to respond to changes in oxygen availability, making it a key cellular oxygen sensing enzyme (3). *Toxoplasma* expresses two oxygen-sensing prolyl hydroxylases named PHYa and PHYb that are important for growth at low and high O_2_, respectively, and appear to act in response to changes in oxygen availability by altering the parasite proteome (4–6).

Besides oxygen, *Toxoplasma* faces diverse nutrient environments within its host and must adapt its metabolic network in response to nutritional variations. The identification of host hexokinase 2 upregulation as a requirement for parasite growth indicated that alterations in host glucose metabolism as one such mechanism (2). Recent dual metabolomic profiling studies further revealed that *Toxoplasma* reprograms the host’s metabolism, shifting it from mitochondrial oxidative phosphorylation to glycolysis as well as increases in host and parasite pentose phosphate pathway (7). Within the parasite, sedoheptulose bisphosphatase, an enzyme that channels sugar molecules from the glycolytic pathway into the pentose phosphate pathway (7), is also active and provides an alternative pathway to generate ribose. Since most of these assays were performed at ambient oxygen, it remains unclear how *Toxoplasma* adjusts its metabolome under low oxygen conditions to fulfill its nutritional needs.

Heme is a vital metabolite in nearly all organisms and is involved in many fundamental metabolic processes, such as cytochrome formation, cellular redox defense, and oxygen sensing and transport. In most eukaryotic cells, the heme biosynthetic pathway comprises eight reactions distributed across two subcellular locations, the cytoplasm and mitochondria (8, 9). Our previous work along with other two research articles have revealed that *Toxoplasma* parasites possess a complete and functional heme biosynthesis pathway (10–12). However, four reactions occur within the apicoplast, a vestigial plastid believed to have originated from endosymbiosis (13). *Toxoplasma* parasites heavily depend on their own heme production for intracellular growth and acute virulence (10–12). Within the heme biosynthesis pathway of *Toxoplasma*, only the antepenultimate reaction takes place in the cytoplasm, producing protoporphyrinogen IX (PROTOgen IX) from coproporphyrinogen III (COPROgen III) catalyzed by coproporphyrinogen III oxidase (CPOX). CPOX utilizes oxygen as an electron acceptor to decarboxylate two propionic acid groups on the A and B pyrrole rings of COPROgen III, ultimately yielding PROTOgen IX (14). In contrast, certain prokaryotic cells like *E. coli* and *Salmonella* encode coproporphyrinogen dehydrogenase, which utilizes radical S-adenosylmethionine (SAM) as an electron acceptor to convert COPROgen III to PROTOgen IX (15, 16). The possession of dual enzymes in these bacteria enables flexible adaptation to various growth conditions.

By analyzing the genome of *Toxoplasma* parasite, an ortholog of bacterial coproporphyrinogen dehydrogenase-like protein, a radical S-adenosyl-L-methionine (SAM) enzyme has been identified and named TgCPDH (TGGT1_288640). Deletion of *TgCPOX* leads to a Δ*cpox* mutant with significant growth defects (10), suggesting that the parasites possess a bypass pathway for heme production, possibly utilizing TgCPDH as an alternative enzyme. A previous study localized TgCPDH to the parasite’s mitochondrion via endogenous gene tagging, and its deletion did not result in the loss of virulence in Type II *Toxoplasma* parasites (12).

Here, we conduct a transgenera complementation experiment of expressing exogenous TgCPDH in a heme synthesis-deficient *Salmonella* mutant (16) to prove that TgCPDH is a functional enzyme in heme biosynthesis. Our findings also revealed that TgCPDH expression increased in response to hypoxia, but its deletion or overexpression in heme-deficient Δ*cpox* parasites did not alter their growth, suggesting that TgCPDH is not directly involved in heme production in *Toxoplasma*. In summary, our research identified a hypoxia-responsive radical SAM enzyme in *Toxoplasma* with coproporphyrinogen dehydrogenase activity, marking the first observation of such an enzyme in a eukaryotic organism.

## RESULTS

### 1. *Toxoplasma* encodes an ortholog of the oxygen-independent heme biosynthesis enzyme, coproporphyrinogen dehydrogenase (CPDH)

Some prokaryotes employ two distinct strategies to convert COPROgen III into PROTOgen IX during the heme biosynthetic pathway. They can utilize either the oxygen-dependent coproporphyrinogen oxidase (HemF) (17, 18) or the oxygen-independent coproporphyrinogen dehydrogenase (HemN) (16, 19, 20) to complete the antepenultimate step in this process. HemF orthologs are prevalent in both eukaryotic and prokaryotic cells. In mammalian cells, this ortholog is known as coproporphyrinogen III oxidase (CPOX) (8). In contrast, HemN orthologs have not been identified in eukaryotic cells. While exploring the genome of *Toxoplasma*, we discovered a radical S-adenosyl-L-methionine (SAM) enzyme, TGGT1_288640, which exhibited similarity to *E. coli* HemN (EcHemN) (**Fig. 1A**). A pairwise alignment revealed 10.4% sequence identity and 25.4% residue similarity using the BLOSUM 45 matrix (**Fig. 1A**). Moreover, according to the crystal structure of *E. coli* HemN, we found that four crucial residues within the catalytic site and a CXXXCXXC motif were conserved in the *Toxoplasma* ortholog (**Fig. 1A**). Therefore, we named this enzyme TgCPDH throughout this paper. TgCPDH possesses a long nonhomologous C-terminal tail relative to EcHemN (**Fig. 1A**). Furthermore, we superimposed the alphafold-predicted structure of TgCPDH (21) with the solved crystal structure of HemN (19) and observed that both proteins exhibited a highly similar core structure, consisting of a group of α-helices and β-sheets (**Fig. 1B**), indicating that TgCPDH structurally resembles *E. coli* coproporphyrinogen dehydrogenase.

**Figure 1.**
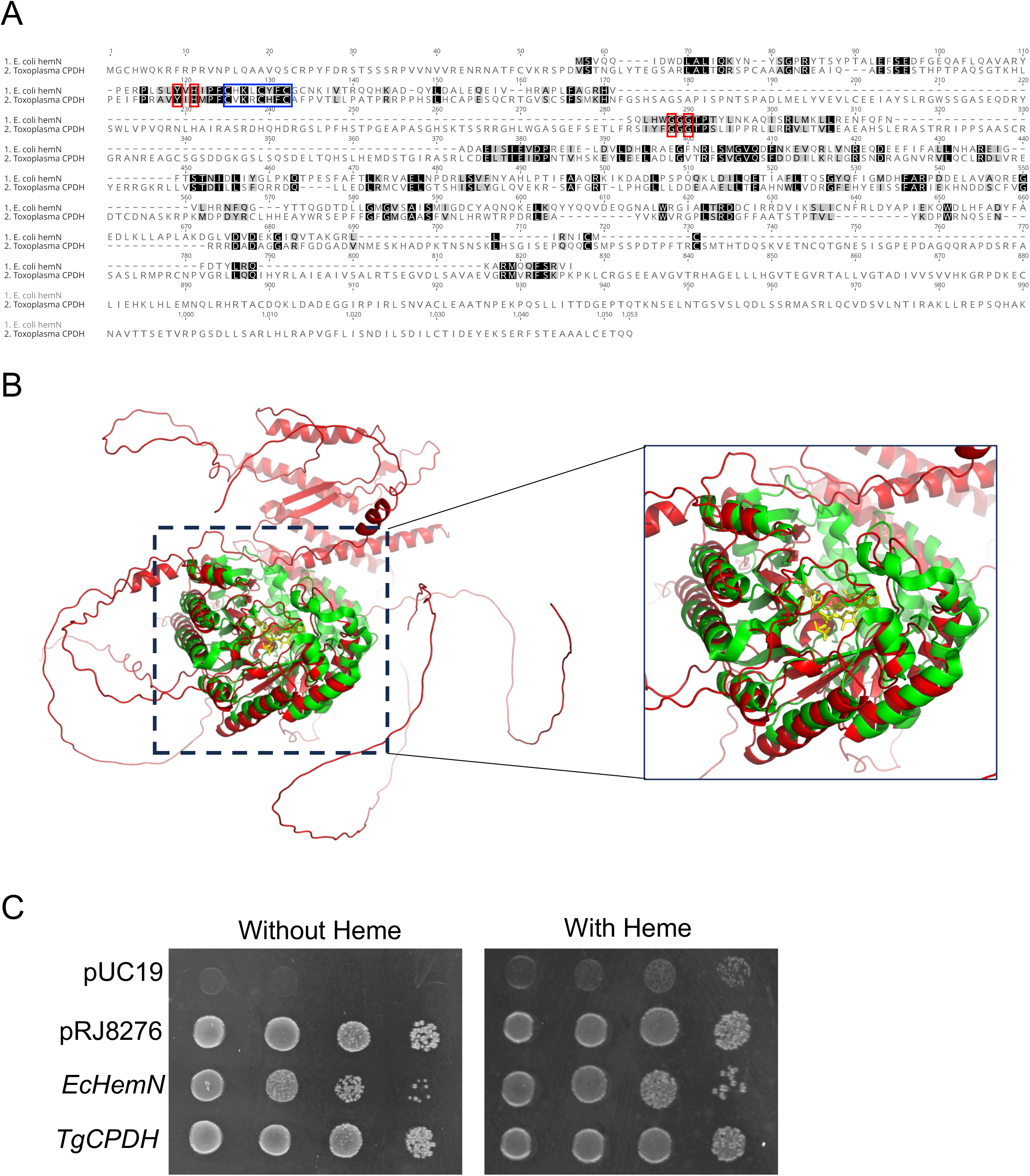
*Toxoplasma* encoded a coproporphyrinogen dehydrogenase-like protein. (A) Comparison of the primary structures of *E. coli* HemN (EcHemN) and *Toxoplasma* CPDH (TgCPDH) through pairwise alignment using ClustalW. The catalytic residues, as determined from the crystal structure of EcHemN, were highlighted with red boxes, while the iron-sulfur cluster binding motif, CXXXCXXC, was enclosed in a blue box. The UniProt IDs for *EcHemN* and *TgCPDH* proteins are P32131 and S7W4N9, respectively. (B) Molecular docking analysis of EcHemN and TgCPDH. The crystal structure of EcHemN was obtained from the RCSB PDB database (PDB ID: 1OLT), while the predicted structure of TgCPDH was generated using the AlphaFold algorithm. Structural alignment was performed using PyMol, with EcHemN and TgCPDH depicted in green and red cartoon backbones, respectively. Two SAM molecules were shown in yellow. (C) Transgenera complementation of *TgCPDH* in the heme-auxotrophic *Salmonella* TE3006 strain restored the growth of transformed bacteria in the medium lacking exogenous heme. The plasmid pRJ8276, encoding *Bradyrhizobium japonicum* HemN2, served as a positive control. Furthermore, *EcHemN* and *TgCPDH* were cloned into pUC19 under the lac promoter and then introduced into *Salmonella* TE3006. An empty pUC19 plasmid was included as a negative control.

To assess whether TgCPDH functions as a coproporphyrinogen dehydrogenase, we PCR-amplified the region with high similarity to EcHemN based on the alignment of primary sequences. Then, we cloned this region under a lac promoter in the pUC19 vector and introduced it into a heme auxotrophic *Salmonella* strain, TE3006, for transgenera complementation. The *Salmonella* TE3006 strain lacks both *hemF* and *hemN* and cannot grow in a medium without heme supplementation (16). We introduced an empty pUC19, pRJ8276 plasmid encoding *Bradyrhizobium japonicum hemN2* (22), and *E. coli hemN*-encoding plasmid individually as negative and positive controls. Our results demonstrated that TgCPDH successfully restored the growth of the *Salmonella* TE3006 strain in a medium lacking exogenous heme (**Fig. 1C**), indicating that TgCPDH indeed exhibits coproporphyrinogen dehydrogenase activity.

### 2. The expression of TgCPDH responded to hypoxic conditions, but it did not facilitate parasite growth under low oxygen levels

Given that TgCPDH exhibited structural similarity to HemN and functionally complemented the growth of heme auxotrophic *Salmonella*, it is speculated that TgCPDH is responsive to low oxygen conditions. To investigate this, we cultured the RHΔ*ku80::TgCPDH^myc^* strain under both ambient and low oxygen conditions (21% and 3%, respectively) and assessed its protein expression through immunoblotting. Our analysis revealed that the translation level of TgCPDH increased under hypoxia compared to that under ambient oxygen (**Fig. 2A**). Subsequently, we hypothesized that TgCPDH can support parasite growth under low oxygen conditions. To test this, we genetically deleted *TgCPDH* in RHΔ*ku80::nLuc*, resulting in a Δ*cpdh::nLuc* strain (**Fig. S1**). Both strains were inoculated into human foreskin fibroblasts (HFFs) for a plaque assay under 21% and 3% oxygen conditions. The plaques were allowed to grow for 7 days before staining with crystal violet and quantification using optical microscopy. The plaques formed by RHΔ*ku80::nLuc* parasites under 3% oxygen were significantly smaller than those formed under 21% oxygen conditions. However, the plaques of Δ*cpdh::nLuc* under 3% oxygen showed comparable sizes to those formed under ambient oxygen (**Fig. 2B**), suggesting that TgCPDH is not involved in parasite growth in hypoxic conditions. Additionally, we investigated the role of TgCPDH in the parasite’s acute virulence. During *in vivo* dissemination, tissue oxygen concentrations are generally lower than those in *in vitro* tissue culture (23). We subcutaneously inoculated individual mice with 100 RHΔ*ku80::nLuc* parasites and either 100 or 1000 Δ*cpdh::nLuc* parasites and monitored mouse mortality daily. In comparison to the parental strain, Δ*cpdh::nLuc* parasites did not exhibit attenuated virulence. Mice infected with both low and high inoculum doses of Δ*cpdh::nLuc* parasites succumbed to infection within 12-13 days post-infection, a timeframe similar to that of RHΔ*ku80::nLuc*-infected mice (**Fig. 2C**). Overall, these results indicate that the expression of TgCPDH was altered in the parasites under different oxygen conditions, but TgCPDH did not play a role in parasite growth under hypoxic conditions, as observed in both *in vitro* and *in vivo* infections.

**Figure 2.**
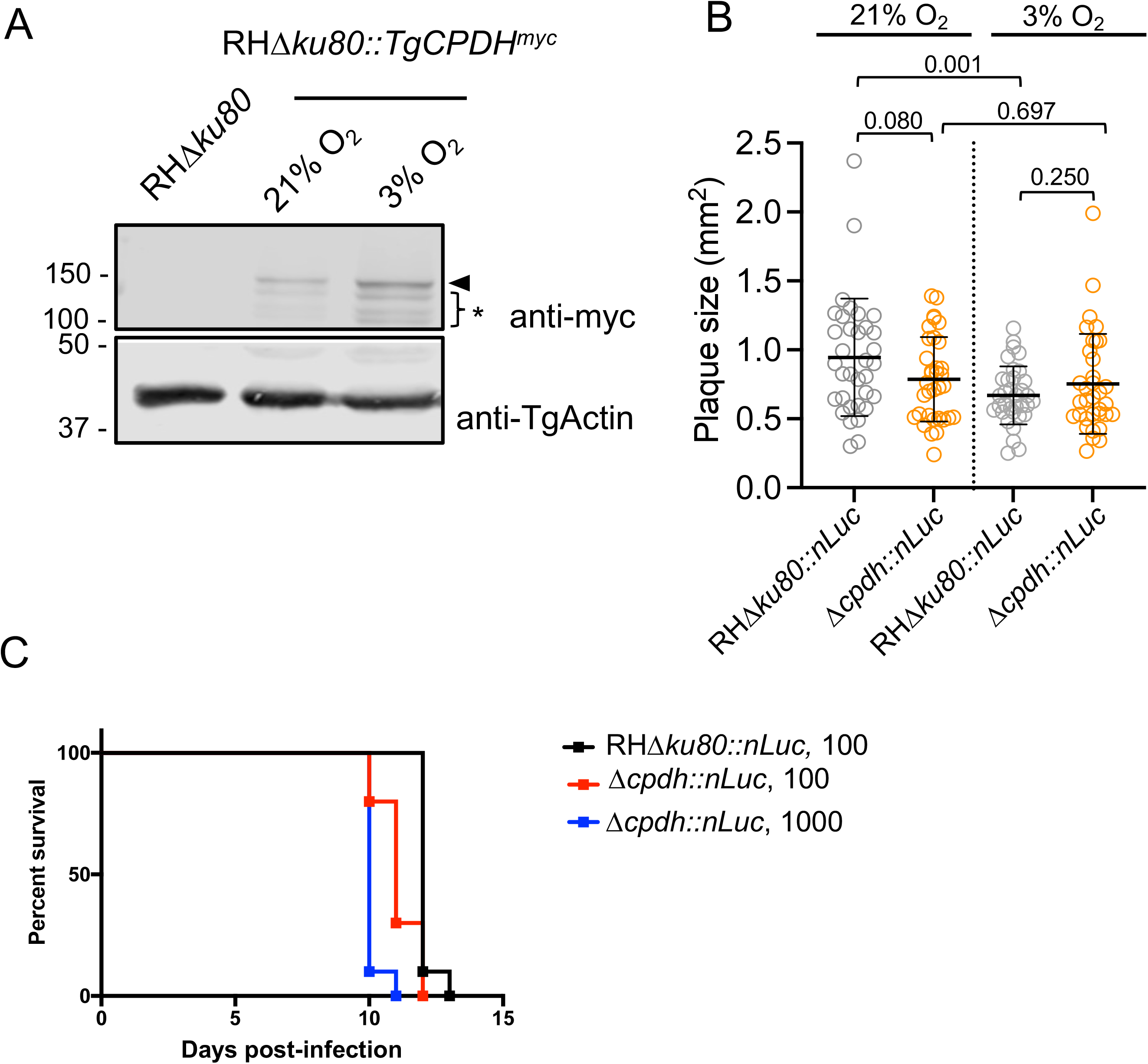
*Toxoplasma* increased TgCPDH expression in response to hypoxia but did not require it for parasite growth under low oxygen conditions. (A) Immunoblotting analysis revealed an elevation in TgCPDH expression in *Toxoplasma* parasites when cultured under 3% O_2_ conditions compared to those grown under ambient O_2_ levels. Lysates were probed with anti-myc antibody for TgCPDH detection. *Toxoplasma* Actin was also probed as a loading control. The full-length TgCPDH band was indicated by an arrowhead, and a series of smaller bands were denoted with an asterisk, possibly representing degradation products. (B) *TgCPDH*-deficient parasites were grown under both ambient and 3% O_2_ conditions for 7 days to evaluate the role of TgCPDH in parasite growth via plaque assay. While wildtype parasites (RHΔ*ku80::nLuc*) formed smaller plaques under 3% O_2_ compared to 21% O_2_, Δ*cpdh::nLuc* parasites did not exhibit an increased growth reduction under 3% O_2_ compared to ambient O_2_ conditions. Statistical significance was assessed using a two-tailed unpaired Student’s *t*-test, with *p*-values indicated in the plot. (C) TgCPDH was dispensable for acute virulence of Type I *Toxoplasma* parasites. CD-1 mice received subcutaneous injections of 100 RHΔ*ku80::nLuc*, 100 or 1000 Δ*cpdh::nLuc* parasites, and mouse mortality was continuously monitored and recorded daily.

### 3. TgCPDH did not facilitate the growth of heme-deficient *Toxoplasma* tachyzoites

Due to the successful transgenera complementation of TgCPDH in the heme auxotrophic *Salmonella* strain, we expected that TgCPDH could potentially compensate for the absence of TgCPOX and aid in completing heme production since they both can convert COPROgen III to PROTOgen IX. To investigate the transcription and translation levels of TgCPDH in heme-deficient parasites, we employed quantitative RT-PCR and immunoblotting to assess the mRNA and protein abundances of TgCPDH in wildtype RHΔ*ku80*, Δ*cpox*, and Δ*cpoxCPOX* parasites, respectively. Under standard growth conditions of 21% oxygen, Δ*cpox* parasites exhibited approximately a 2.5-fold higher transcript level of TgCPDH compared to wildtype parasites (**Fig. 3A**). For protein quantification, initially, we attempted to endogenously tag TgCPDH with a C-terminal 3x-HA epitope tag. However, the signals for both immunoblotting and immunofluorescence proved to be very weak. Consequently, we opted to genetically insert a Spaghetti-Monster 10xmyc (smGFP-myc) epitope tag (24, 25) at the C-terminus of TgCPDH using CRISPR-Cas9-mediated genome manipulation. The predicted molecular weight (MW) of TgCPDH is approximately 112 kDa, while that from the smGFP-myc tag is approximately 40 kDa. As anticipated, the apparent MW for TgCPDH-smGFP-myc was around 150 kDa (**Fig. 3B**). We also probed the lysates with anti-TgSAG1 antibody as a loading control for normalization. The quantitative analysis revealed that the protein level of TgCPDH in Δ*cpox* was comparable to that in the wildtype and Δ*cpoxCPOX* strains. We further conducted an immunofluorescence analysis to assess if TgCPDH alter its subcellular localization in Δ*cpox*. Our data demonstrated that TgCPDH remained within the mitochondria in all three strains (**Fig. 3C**), and the fluorescence intensities of anti-myc signals were comparable, corroborating our immunoblotting quantification. To investigate whether TgCPDH played a crucial role in the growth of Δ*cpox* mutant, we genetically deleted the *TgCPDH* gene in the Δ*cpox::nLuc* strain that had been generated in a prior study, resulting in a Δ*cpoxΔcpdh::nLuc* strain (**Fig. S1**). The double knockout strain remained viable under ambient oxygen conditions. We quantified its growth through a plaque assay alongside Δ*cpox::nLuc* and Δ*cpoxCPOX::nLuc* strains under both 21% and 3% oxygen conditions. Remarkably, we did not observe a significant difference in parasite growth between Δ*cpoxΔcpdh::nLuc* and the other strains under regular and hypoxic conditions (**Fig. 3D**). In conclusion, our findings suggest that although the parasites increase transcript levels of TgCPDH in the Δ*cpox* mutant, TgCPDH does not significantly contribute to heme biosynthesis within Δ*cpox* parasites for their intracellular growth.

**Figure 3.**
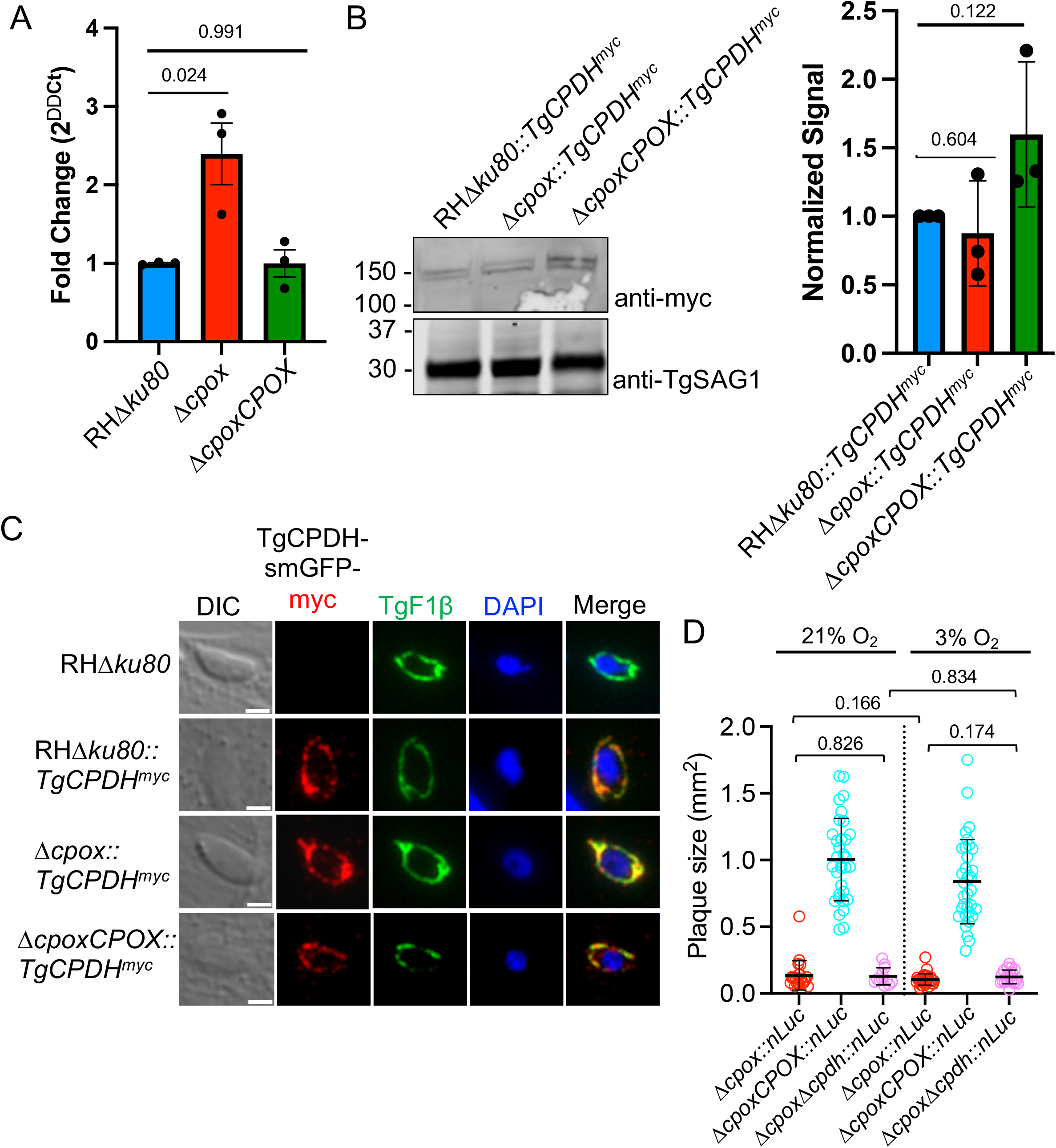
Transcript levels of TgCPDH were increased in the Δ*cpox* mutant, but it was not involved in its intracellular growth. (A) Quantitative RT-PCR analysis revealed approximately a 2.5-fold increase in *TgCPDH* transcript levels in Δ*cpox* compared to RHΔ*ku80* and Δ*cpoxCPOX* parasites. *Toxoplasma* RNA polymerase II was used as a housekeeping gene for normalization. (B) Quantitative immunoblotting analysis did not detect the elevated expression of TgCPDH in Δ*cpox* mutant. TgSAG1 was used as a loading control. (C) Immunofluorescence assay demonstrated that TgCPDH remained localized within the mitochondria of Δ*cpox* parasites. TgF1β was used as a marker of *Toxoplasma* mitochondrion. Scale bar: 2 µm. (D) Deletion of *TgCPDH* in the Δ*cpox* mutant did not result in a reduction in parasite growth under both ambient and 3% O_2_ conditions, as determined by plaque assay. Plaque areas were measured and presented as mean sizes ± standard deviations. Statistical significance was calculated using a two-tailed unpaired Student’s *t*-test, with *p*-values marked in the plot.

### 4. The overexpression of TgCPDH did not result in an increased parasite growth rate in heme-deficient parasites

The 2.5-fold increase in TgCPDH mRNA levels observed in Δ*cpox* did not translate into a significant elevation of TgCPDH protein expression. This suggests that heme metabolism may not be substantially improved in Δ*cpox*, thereby continuing to limit parasite growth. To address this challenge, we conducted an overexpression experiment by placing *TgCPDH* under the control of a *Toxoplasma* tubulin promoter and adding a C-terminal 3xHA epitope for immunodetection. These plasmids were introduced into RH*Δku80::nLuc* and Δ*cpox::nLuc* strains, resulting in the creation of RHΔ*ku80::nLuc/pTub-TgCPDH^HA^*and Δ*cpox::nLuc/pTub-TgCPDH^HA^* strains. First, we assessed the mRNA levels of TgCPDH in the TgCPDH-overexpression strains using quantitative RT-PCR. The analysis revealed that the mRNA levels were dramatically elevated compared to strains not overexpressing TgCPDH and the transcription levels were similar in both RHΔ*ku80::nLuc/pTub-TgCPDH^HA^*and Δ*cpox::nLuc/pTub-TgCPDH^HA^* (**Fig. 4A**). Second, we examined the protein levels of the overexpressed TgCPDH in both strains. Similarly, the protein levels of TgCPDH were comparable when overexpressed in both RHΔ*ku80* and Δ*cpox* (**Fig. 4B**). To ensure that overexpressed TgCPDH maintained its subcellular localization, we conducted immunofluorescence staining using anti-HA antibodies along with anti-TgF1β for *Toxoplasma* mitochondrion recognition. The analysis demonstrated that overexpressed TgCPDH still remained localized to the mitochondria (**Fig. 4C**). We monitored parasite growth by measuring luciferase activity in these strains, as they expressed nanoLuciferase. The strains were cultured in confluent HFFs, and parasite growth was assessed every 24 hrs over a total period of 96 hrs. We did not observe an enhanced parasite growth in Δ*cpox::nLuc/pTub-TgCPDH^HA^*relative to Δ*cpox::nLuc*. Similarly, overexpression of TgCPDH in RHΔ*ku80* did not result in improved growth either. In summary, although TgCPDH was effectively overexpressed in *Toxoplasma* parasites, such overexpression did not lead to an increase in the growth of heme-deficient parasites under standard ambient oxygen growth conditions (**Fig. 4D**).

**Figure 4.**
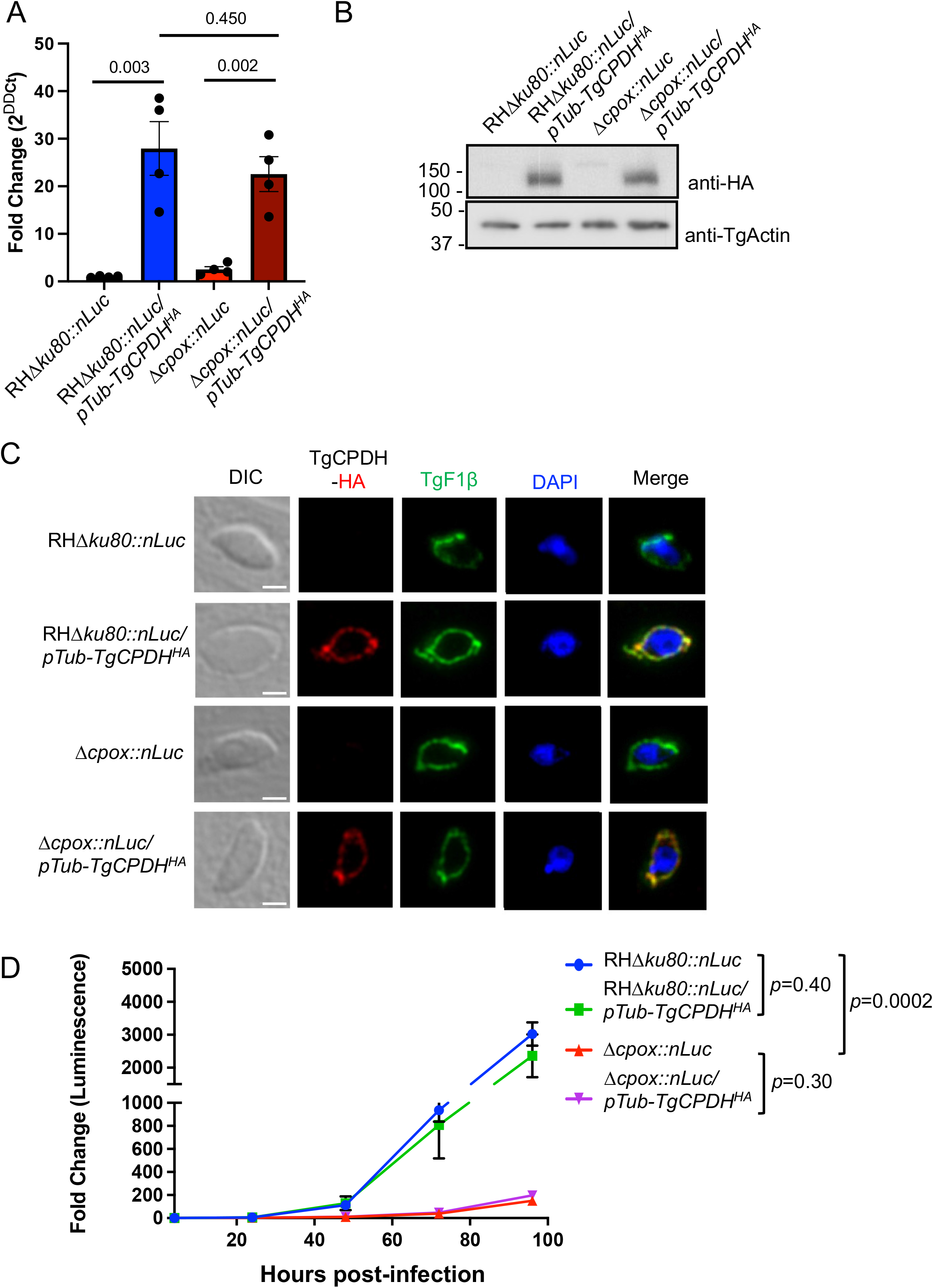
Overexpression of TgCPDH did not enhance the growth rate of Δ*cpox*. TgCPDH overexpression was achieved by placing exogenous TgCPDH under the control of a *Toxoplasma* tubulin promoter, and a 3xHA tag was added to its C-terminus for immunodetection. (A) Quantitative RT-PCR analysis demonstrated a significant increase in TgCPDH transcript levels in the strains overexpressing *TgCPDH*. *Toxoplasma* RNA polymerase II served as a housekeeping gene for normalization. (B) Immunoblotting analysis revealed a comparable expression level of exogenous overexpressed TgCPDH in both RHΔ*ku80* and Δ*cpox* parasites. TgActin served as a loading control. (C) Immunofluorescence assay confirmed that overexpressed TgCPDH retained its localization within the mitochondrion of *Toxoplasma*. TgF1β was included as a marker of *Toxoplasma* mitochondria. Scale bar: 2 µm. (D) A luciferase-based growth analysis did not detect significant improvement in parasite growth in the Δ*cpox* mutant overexpressing TgCPDH compared to Δ*cpox* alone. The assay was conducted in triplicate, and the statistical significance of the growth rates at 96 hrs post-infection was determined using a two-tailed unpaired Student’s *t*-test, with *p*-values indicated in the plot.

## DISCUSSION

Over 600 members of the radical SAM protein superfamily have been identified through bioinformatic searches (26). This group of proteins utilizes an unconventional [4Fe-4S]^+^ cluster as a cofactor to catalyze a reductive cleavage of S-adenosyl-L-methionine (SAM), generating a radical species, typically a 5-deoxyadenosyl radical (5’-dAdo), which abstracts hydrogen atoms of the substrates they catalyze (27, 28). These radical-based enzymatic reactions are believed to represent an ancient and conserved mechanistic approach to handling challenging chemical reactions (26). Radical SAM proteins catalyze a wide range of reactions, including the anaerobic oxidations of COPROgen III into PROTOgen IX (16, 20, 22), a critical step within the heme biosynthetic pathway. So far, this reaction has only been observed in some bacteria and is catalyzed by oxygen-independent HemN/HemZ proteins (16, 20, 22, 29). However, a recent examination of the *Toxoplasma* genome has revealed the presence of an ortholog of HemN, therefore named *Toxoplasma* coproporphyrinogen dehydrogenase (TgCPDH) (12). Notably, a HemZ ortholog was not found in the *Toxoplasma* genome. Through primary structure alignment, it is evident that TgCPDH possesses all four catalytic residues found in *E. coli* HemN, specifically Y^56^, H^58^, G^111^, and G^113^, and additionally features a CXXXCXXC motif for [4Fe-4S]^+^ cluster binding (19, 20, 30, 31). Our transgenera complementation of TgCPDH in a *Salmonella* mutant deficient in converting COPROgen III to PROTOgen IX demonstrated that TgCPDH exhibits coproporphyrinogen dehydrogenase activity. Notably, bacterial CPDH enzymes encode a fourth Cys preceded by the [4Fe-4S]^+^ binding site to form a CXXXCXXCXC motif, with the first three cysteines being responsible for iron-sulfur cluster binding, while the fourth plays a role in catalysis (20, 22, 32). Some proteins previously classified as HemN have been reclassified as HemW due to the absence of this critical cysteine. As anticipated, HemW cannot rescue the growth of heme auxotrophic *Salmonella* mutants (32). Instead, it acts as a heme chaperone, facilitating heme trafficking and insertion into hemoproteins (32). In the case of TgCPDH, only the CXXXCXXC motif is observed, but several cysteine residues follow this motif (**Fig. 1A**). These additional cysteines may be structurally proximate to the [4Fe-4S]^+^-binding motif and potentially mediate enzymatic catalysis. It is also plausible that TgCPDH functions as a heme chaperone, aiding in the transport and conjugation of heme into mitochondrial hemoproteins, given its mitochondrial localization. To investigate this possibility further, a series of biochemical assays assessing mitochondrial respiration and mitochondrial membrane potentials can be conducted in both wildtype and Δ*cpdh* parasites, which will provide insights into the potential role of TgCPDH in the parasite’s mitochondrial health.

The immunofluorescence assay has confirmed the mitochondrial localization of TgCPDH, but its exact sub-mitochondrial location remains unknown. Notably, TgCPOX catalyzes the conversion of COPROgen III into PROTOgen IX in the cytosol of the parasites (10). If TgCPDH is located in the mitochondrial intermembrane space or matrix, COPROgen III must cross the mitochondrial membrane(s) via a transporter before undergoing catalysis. In mammalian cells, the ABCB6 transporter has been identified as mediating the import of heme precursors across mitochondrial membranes (33), facilitating the final steps of heme biosynthesis from the cytosol to the mitochondria. *Toxoplasma* contains an ortholog of ABCB6 (TGGT1_269000), but its function requires further investigation.

Despite our findings indicating that TgCPDH exhibits CPDH function via transgenera complementation, the deletion of this ortholog in heme-deficient *Toxoplasma* Δ*cpox* parasites did not result in arrested growth. This finding weakens the hypothesis that TgCPDH is directly involved in heme biosynthesis, although its contribution to heme production cannot be completely ruled out. Since the Δ*cpox* mutant remains viable in tissue culture, it is possible that alternative enzymes may facilitate endogenous heme production or that the parasite may acquire heme or heme biosynthetic intermediates from the host. Previous research has shown that the deletion of the last enzyme, TgFECH, within the heme biosynthetic pathway, is lethal (10, 12). Additionally, the addition of exogenous heme in the medium did not enhance the growth of heme-deficient mutants (10, 12). These findings suggest that *Toxoplasma* parasites are not able to acquire heme from host cells or cannot obtain sufficient amounts to support parasite growth. A previous study proposed that parasites can scavenge PROTO IX or PROTOgen IX, which are the products catalyzed by TgPPO or TgCPOX, respectively, from host cells to boost heme production (12). Notably, PROTOgen IX can be autonomously oxidized into PROTO IX, and the primary difference between PROTO IX and intact heme is the absence of a ferrous ion. Therefore, it remains unclear how PROTO IX can be transported into the mitochondria rather than the intact heme molecule. Furthermore, the observation that deletion mutations in *TgALAS* and *TgUROD* mutants, which respectively encode the first and fifth enzyme within the pathway, grew poorly or died in standard D10% medium (10, 11), weakens the speculation that *Toxoplasma* can scavenge PROTO IX from its host. Future investigations involving isotope-labeled heme or heme intermediates may help trace the fate of host heme or its precursors in the parasites. Furthermore, a study has shown that a HemY protein from *Bacillus subtilis* can catalyze both CPOX and PPO reactions (34). Hence, it is possible that TgPPO can perform the function of TgCPOX, albeit with lower efficacy, potentially explaining the viability of the Δ*cpox* mutant. Further experiments testing whether TgPPO can rescue the growth of CPOX-deletion *Saccharomyces cerevisiae* in a medium lacking heme may help address this question.

Our study has revealed that TgCPDH expression is responsive to low oxygen conditions. In mammalian cells, hypoxia can lead to an accumulation of reactive oxygen species (ROS) in the mitochondria, resulting in cellular damage (35). Consequently, cells must regulate mitochondrial activity to control ROS production (36). Interestingly, our Δ*cpdh* mutant did not exhibit more severe growth defects under hypoxia conditions; instead, it grew at a similar rate to ambient oxygen conditions. Quantifying ROS levels in the Δ*cpdh* mutant under both ambient and hypoxia conditions may help determine if CPDH expression levels correlate with ROS levels in the parasites. It is possible that a compensation mechanism operates in the Δ*cpdh* mutant to offset its loss. Future transcriptomics and proteomics comparisons between wildtype and *TgCPDH*-deficient parasites may shed light on this compensation mechanism.

## MATERIALS AND METHODS

### Ethical statement

CD-1 outbred mice were subcutaneously individually injected with 100 RHΔ*ku80::nLuc*, 100 or 1,000 Δ*cpdh::nLuc* parasites prepared in 1X Phosphate-Buffered Saline (PBS). Mice were monitored post-infection for adverse symptoms for a course of 30 days. Mice that showed over 20% drop in initial weight were euthanized by CO_2_ in compliance with the protocol approved by Clemson University’s Institutional Animal Care and Use Committee (Animal Welfare Assurance A3737-01, protocol number 2019-035). The survival rate was plotted, and statistical analysis was calculated using the Log-rank (Mantel-Cox) test.

### Chemicals and reagents

The chemicals used in this study were of analytical grade and were acquired from VWR, unless specified otherwise. All oligonucleotide primers used in this work, as listed in Table S1, were obtained from Eurofins.

### Host cell and parasite culture

Human foreskin fibroblasts (HFFs) were provided by the American Type Culture Collection (ATCC, catalog #: SCRC-1041). HFFs were grown in D10% growth medium consisting of DMEM (Dulbeccos’s Modified Eagle Medium), 4.5 g/L glucose, 10% Cosmic Calf Serum (Hyclone, SH30087.03), 10 mM HEPES, 4 mM glutamine, and 10 mM Pen/Strep. Host cells and parasites were incubated at 37°C with 5% CO_2_ and either 21% or 3% O_2_. All *Toxoplasma* strains used for this study were maintained by two-day serial passage in HFF cells supplemented with D10% media before use in all assays.

### Complementation of *Toxoplasma* coproporphyrinogen dehydrogenase-like ortholog into ***Salmonella* TE3006**

#### (1) Preparation of competent *Salmonella* TE3006 cells

The *Salmonella* TE3006 strain, lacking both the oxygen-dependent coproporphyrinogen III oxidase (HemF) and the oxygen-independent coproporphyrinogen dehydrogenase (HemN), was generously provided by Dr. Hans-Martin Fischer from ETH Zürich. Competent *Salmonella* TE3006 cells were prepared using the Zymo Mix and Go kit (Zymo Research), following the provided instructions. In brief, the competent bacteria were cultured in Luria-Bertani broth (LB) supplemented with 15 µg/mL hemin, tetracycline, and kanamycin until they reached an optical density of 600 nm (OD_600_) of 0.6. Subsequently, the cells were centrifuged at 3,000 x g for 10 min at 4°C and washed twice with a wash buffer to remove residual hemin. The competent cells were then resuspended in a competent buffer and stored at -80°C for later use in the transformation process.

#### (2) Construction of plasmids expressing *E. coli HemN* or *Toxoplasma TgCPDH*

The *E. coli HemN* (*EcHemN*) gene was PCR-amplified from *E. coli* genomic DNA and inserted into the pUC19 plasmid using BamHI and EcoRI restriction sites. Based on the high homologous region between *TgCPDH* and *EcHemN* from primary structure alignment, we isolated the DNA sequence corresponding to a partial region of TgCPDH, spanning from residue 38 to 790, from a *Toxoplasma* cDNA library, which contains the conserved catalytic residues and motifs. This sequence was subsequently cloned into the pUC19 plasmid through Gibson DNA assembly. Additionally, we incorporated a 3xHA epitope tag at the C-termini of both EcHemN and the truncated TgCPDH.

#### (3) Transformation of *Salmonella* TE3006 Cells

*Salmonella* transformations were carried out by combining 100 µL of competent cells with 2 µL of plasmid DNA (400-700 ng/µL) in a culture tube, followed by incubation on ice for 30 min. The mixture was then supplemented with 1 mL of SOC medium containing 20 mM glucose and shaken at 225 rpm at 37°C for 1 hr. Subsequently, the bacteria were centrifuged at 3,000 x g at room temperature for 10 min, and the resulting pellet was resuspended in 200 µL of SOC medium with glucose. The suspension was then spread onto LB plates containing ampicillin, kanamycin, and tetracycline, with and without 15 µg/mL hemin. These plates were incubated at 30 °C overnight to allow for the formation of single colonies.

#### (4) Assessment of growth in transformed *Salmonella* TE3006 cells

The individual clones of *Salmonella* TE3006 strains transformed with plasmids expressing *Bradyrhizobium japonicum HemN2* generously provided by Dr. Hans-Martin Fischer from ETH Zürich, *E. coli HemN*, or *Toxoplasma TgCPDH* were cultured overnight in the medium supplemented with ampicillin, kanamycin, and tetracycline, both with and without 15 µg/mL hemin. The cultures were then diluted in 1x PBS to achieve an OD_600_ of 0.8, with three additional 10-fold serial dilutions. Two microliters of the diluted bacterial suspensions were spotted onto LB plates supplemented with ampicillin, kanamycin, tetracycline, and 10 mM IPTG, with and without 15 µg/mL hemin. After drying for 3 hrs, the plates were incubated at 30 °C for 3 days before imaging.

### Generation of transgenic *Toxoplasma* strains (Table S2)

#### (1) Generation of *Δcpdh::nLuc* and Δ*cpox*Δ*cpdh::nLuc* strains

To create the *Δcpdh::nLuc* and Δ*cpox*Δ*cpdh::nLuc* strains, following established protocols (10, 37), we employed CRISPR-based genome modification techniques to delete the *TgCPDH* gene (TGGT1_288640) in RHΔ*ku80*Δ*hxg::nLuc* and *Δcpox::nLuc*. First, we designed a guide RNA targeting the 3’-end of the *TgCPDH* gene, as previously described (10, 37). Second, we used primers carrying 50-base pair homologous regions flanking the start and stop codons of *TgCPDH* to amplify a pyrimethamine resistance cassette. After electrophoresis and gel extraction, we mixed the repair template with the guide RNA and introduced it into filter-purified *Toxoplasma* parasites suspended in Cytomix buffer through electroporation, as detailed in previous studies (10, 37). Following pyrimethamine selection, we cloned out the mutant parasites in 96-well plates pre-seeded with HFFs. The correct mutant clones were screened by PCR to confirm the correct integration of 5’- and 3’-ARMs and the removal of the *TgCPDH* coding sequence.

#### (2) Generation of spaghetti-monster-myc tag (smGFP-myc)-labeled TgCPDH strains

To label the TgCPDH gene with a smGFP-myc epitope tag, we genetically inserted this tag at the C-terminus of the *TgCPDH* gene in RH*Δku80*, Δ*cpox*, and Δ*cpoxCPOX* parasites using the CRISPR technique. Similar to the previous procedure, 50-base pair homologous regions flanking the stop codon of TgCPDH were placed at the 5’- and 3’-ends of the smGFP-myc epitope tag and a chloramphenicol resistance cassette via PCR. We combined this repair template with a guide RNA targeting the 3’-end of the *TgCPDH* gene and introduced them into *Toxoplasma* parasites via electroporation, as outlined above. After transfection, we selected the parasites using chloramphenicol and subsequently isolated individual clones. Correct gene tagging was screened through PCR and immunoblotting.

#### (3) Generation of *Toxoplasma* Strains Overexpressing TgCPDH

We PCR-amplified the coding sequence of *TgCPDH* from a *Toxoplasma* cDNA library and constructed it into a plasmid containing a pyrimethamine resistance cassette. The *TgCPDH* gene was driven by a *Toxoplasma* tubulin promoter (pTub) for its overexpression, and a 3xHA epitope was added to the C-terminus of TgCPDH for immunodetection. The resulting plasmids were introduced into RHΔ*ku80::nLuc* and Δ*cpox::nLuc* parasites using standard electroporation parameters. Following drug selection, we cloned the transfected parasites and subsequently screened them via PCR.

### SDS-PAGE and Immunoblotting

Parasites were grown in confluent HFFs in D10% medium, following a 2-day pass regimen before each experiment. Parasites were harvested by syringing infected host cells with a 25-gauge needle and purifying the parasites through a 3 μm filter. Purified parasites were pelleted at 1,000 x g and then resuspended in 1x SDS-PAGE sample buffer (40 mM Tris, pH 6.8, 1% SDS, 5% glycerol, 0.0003% bromophenol blue, and 50 mM DTT) with the addition of 2% (v/v) β-mercaptoethanol. Subsequently, samples were boiled for 10 min before SDS-PAGE. Proteins resolved on SDS-PAGE were semi-dry transferred onto PVDF membranes. A chemiluminescent immunoblotting strategy was used to detect target proteins. Initially, the blots were blocked with 5% non-fat milk in PBS-T (PBS buffer with 0.1% (v/v) Tween-20). Primary antibodies were prepared in 1% non-fat milk dissolved in PBS-T. Secondary antibodies, either goat α-mouse or α-rabbit IgG conjugated with horseradish peroxidase, were subsequently applied. To develop chemiluminescence signals, the blots were exposed to SuperSignal WestPico chemiluminescent substrate (Thermo Fisher). Finally, images were captured using the Azure C600 Imaging System.

### Immunofluorescence microscopy

Filter-purified parasites were introduced into 6-well chamber slides containing confluent monolayer HFFs. Parasites were cultured in D10% medium at 37°C, with an atmosphere of 5% CO_2_ and 21% O_2_, for 4 hrs. Following this initial incubation, non-invaded parasites were washed away, and fresh D10% medium was replaced. Slides were then incubated for an additional 20 hrs to allow the parasites to form parasitophorous vacuoles, each containing 4-8 parasites, before formaldehyde fixation. To stain the intracellular parasites, infected HFFs were permeabilized using a 0.1% TritonX-100 in 1x PBS. Staining was performed using mouse anti-F1β antibodies to target the *Toxoplasma* mitochondrion, along with rabbit anti-HA or rabbit anti-myc antibodies to label TgCPDH. Subsequently, the slides were stained with goat anti-mouse or goat anti-rabbit IgGs conjugated with different fluorescent dyes (Alexa 488 or 594, Thermo Fisher). Observations were made using a Leica DMi8 inverted epifluorescence microscope at a magnification of 1,000x, equipped with a CCD camera. The captured images were then processed using Leica LAS X software.

### Quantitative reverse-transcriptase PCR (qRT PCR)

*Toxoplasma* parasites were cultured in HFFs for a two-day period prior to the extraction of total RNA. Extracellular parasites were subjected to filter-purification and resuspended in ice-cold 1X PBS. Total RNA was then extracted from the parasites using the Direct-zol RNA MiniPrep Plus kit (Zymo Research). Approximately 100 ng of total RNA was used in the detection and quantification of TgCPDH transcripts using the Luna Universal One-Step RT-PCR kit (NEB). Data acquisition was collected using the BioRad CFX96 Touch Real-Time PCR detection system. A double delta cycle threshold (ΔΔCT) analysis was applied to determine the abundance of TgCPDH transcripts relative to those in wildtype parasites, following previously reported procedures (37). *Toxoplasma* RNA Polymerase II was used as a housekeeping gene for normalization.

### Luciferase-based growth assay

Parasites were filter-purified in phenol-red free D10% medium and subsequently inoculated into 96-well white solid-bottom plates with pre-seeded confluent HFFs. Typically, each well was inoculated with 1,500 tachyzoites. However, for strains displaying growth defects, a higher inoculum of 7,500 tachyzoites per well was used to ensure the generation of reliable signals for measurement. After allowing the parasites to invade the host cells for a 4-hour duration, the culture medium was replaced with fresh phenol-red free D10% to eliminate any non-invaded parasites. Bioluminescence signals were then recorded at 24-hr intervals over a total period of 96 hrs, following established protocols (10, 38). The average signal readings at each time point were divided by the average readings at 4 hrs to calculate the fold change of intracellular growth for each strain.

### Plaque assay

Freshly lysed parasites were filter-purified and subsequently resuspended in room temperature D10% medium, resulting in a parasite concentration of 100 parasites per mL. A half milliliter of this diluted parasite suspension was then inoculated into 24-well plates pre-seeded with confluent HFFs. Plates were placed in incubators at 37°C with 5% CO2, subjecting them to both ambient or low oxygen conditions (21% and 3% O2, respectively) for a continuous period of 7 days without disturbance. Each well was stained with 0.002% crystal violet in 70% ethanol for a duration of 5 min, followed by a thorough rinse with water. To quantify the differences in plaque size, 12-35 plaques were captured at 25x magnification using a Leica DMi8 microscope. These plaque areas were then plotted and analyzed using Prism GraphPad software to assess size variations among different strains.

### Molecular Docking Analysis

The 3D crystal structure of *E. coli* HemN (PDB ID: 1OLT) was retrieved from the RCSB Protein Data Bank (19). Additionally, the predicted 3D structure of TgCPDH was obtained from the AlphaFold protein structure database (21). Subsequently, the alignment of TgCPDH with the *E. coli* HemN structure was carried out using Pymol Molecular Graphics 2.0 (Schrödinger LLC in New York, NY, USA).

### Statistical analysis

All statistical analyses were calculated using GraphPad Prism software (Version 8). Detailed methods for individual assays were specified in the figure legends.

## ACKNOWLEDGMENTS

We thank our colleagues, Drs. Fischer Hans-Martin, Dieter Jahn, Peter Bradley, Silvia NJ Moreno, Vern Carruthers, and David Sibley for generously supplying essential reagents for this study. Funding for this research was provided by the National Institutes of Health grant R01AI143707 (awarded to Z.D.), R01AI169849 (awarded to IJB), and R01AI150240 (awarded to IJB). We declare that we have no conflicts of interest concerning the contents of this article.

## SUPPLEMENTAL MATERIALS

**Figure S1. Creation of Δ*cpdh::nLuc* and Δ*cpoxΔcpdh::nLuc Toxoplasma* strains.** Schematic representation of the standard CRISPR-Cas9-based methodology employed for deleting the *TgCPDH* gene in *Toxoplasma*. The successful integration of the pyrimethamine resistance cassette (DHFR) into the *TgCPDH* locus during gene deletion was verified using PCR. The genomic positions of the primers used for PCR are highlighted within the diagram.

**Table S1. Primers used in this work.**

**Table S2. *Toxoplasma* strains used in this work.**

